# An introduction to thermodynamic integration and application to dynamic causal models

**DOI:** 10.1101/2020.12.21.423807

**Authors:** Eduardo A. Aponte, Yu Yao, Sudhir Raman, Stefan Frässle, Jakob Heinzle, Will D. Penny, Klaas E. Stephan

## Abstract

In generative modeling of neuroimaging data, such as dynamic causal modeling (DCM), one typically considers several alternative models, either to determine the most plausible explanation for observed data (Bayesian model selection) or to account for model uncertainty (Bayesian model averaging). Both procedures rest on estimates of the model evidence, a principled trade-off between model accuracy and complexity. In the context of DCM, the log evidence is usually approximated using variational Bayes. Although this approach is highly efficient, it makes distributional assumptions and is vulnerable to local extrema. This paper introduces the use of thermodynamic integration (TI) for Bayesian model selection and averaging in the context of DCM. TI is based on Markov chain Monte Carlo sampling which is asymptotically exact but orders of magnitude slower than variational Bayes. In this paper, we explain the theoretical foundations of TI, covering key concepts such as the free energy and its origins in statistical physics. Our aim is to convey an in-depth understanding of the method starting from its historical origin in statistical physics. In addition, we demonstrate the practical application of TI via a series of examples which serve to guide the user in applying this method. Furthermore, these examples demonstrate that, given an efficient implementation and hardware capable of parallel processing, the challenge of high computational demand can be overcome successfully. The TI implementation presented in this paper is freely available as part of the open source software TAPAS.

**Author summary:** When fitting computational models to data in the setting of Bayesian inference, a user has the choice between two broad classes of algorithms: variational inference and Monte Carlo simulation. While both methods have advantages and drawbacks, variational inference has become standard in the domain of modelling directed brain connectivity due to its computational efficiency, especially when the challenges to select between competing hypotheses that explain the observed data. By contrast, the high computational demand by Monte Carlo methods has so far prevented their widespread use for inference on brain connectivity, despite their capability to overcome some of the shortcomings of variational inference. In this paper, we introduce the user to thermodynamic integration (TI), a Monte Carlo method designed for model fitting and model selection. By covering its foundations and historical origins in statistical physics, we hope to convey an in-depth understanding of TI that goes beyond a purely technical treatment. In addition, we also provide examples for concrete applications, demonstrating that, given an efficient implementation and up-to-date hardware, the challenge of high computational demand can be overcome successfully.

## Introduction

Dynamic causal models (DCMs) are generative models that serve to infer latent neurophysiological processes and circuit properties – e.g., the effective connectivity between neuronal populations – from neuroimaging measurements such as functional magnetic resonance imaging (fMRI) or magneto-/electroencephalography (M/EEG) data (David et al., 2006; Friston, Harrison, & Penny, 2003). As reviewed by Daunizeau, David, and Stephan (2011), DCMs consist of two hierarchically related layers: a set of state equations describing neuronal population activity, and an observation model which links neurophysiological states to observed signals and accounts for measurement noise. Equipped with a prior distribution over model parameters, a DCM specifies a full generative forward model that can be inverted using Bayesian techniques.

In addition to inference on model parameters, an important scientific problem is the comparison of competing hypotheses, for example, different network topologies, which are formalized as different models. Under the Bayesian framework, model comparison is based on the evidence or marginal likelihood of a model. The model evidence corresponds to the denominator (or normalization constant) from Bayes’ theorem and represents the probability of the observed data under a given model. It is a widely used score of model quality that quantifies the trade-off between model fit and complexity (Bishop, 2006; MacKay, 2004).

Unfortunately, in most instances, it is not feasible to derive an analytical expression of the model evidence due to the intractable integrals that arise from the marginalization of the model parameters. While various asymptotical approximations exist, such as the Bayesian Information Criterion (BIC, Schwarz, 1978) and more recently the Widely Applicable Bayesian Information Criterion (WBIC, Watanabe, 2013), variational Bayes under the Laplace approximation (VBL, Friston, Mattout, Trujillo-Barreto, Ashburner, & Penny, 2007) has established itself as the method of choice for DCM, partially because of its computational efficiency. Within the framework of variational Bayes (VB), a lower bound approximation of the log model evidence (LME) is obtained as a byproduct of model inversion: the variational negative free energy (which we refer to as −*FVB* throughout this paper).

While highly efficient, model comparison based on the variational free energy has several potential pitfalls. For example, under the Laplace approximation as used in the context of DCM, there is no guarantee that −*F*_*VB*_ still represents a lower bound of the LME (Wipf & Nagarajan, 2009). Furthermore, VB is commonly performed in combination with a mean field approximation, and the effect of this approximation on the posterior estimates can be difficult to predict (Daunizeau et al., 2011). Finally, in non-linear models, the posterior could become a multimodal density, a condition that aggravates the application of gradient ascent methods regularly used in combination with the Laplace approximation.

For these reasons, Markov Chain Monte Carlo (MCMC) sampling has been explored as an alternative inference technique for DCM (Aponte et al., 2016; Chumbley, Friston, Fearn, & Kiebel, 2007; Penny & Sengupta, 2016; Sengupta, Friston, & Penny, 2015, 2016; Yao & Stephan, 2020). MCMC is particularly attractive for variants of DCMs in which Gaussian assumptions might be less adequate, such as nonlinear DCMs for fMRI (Stephan et al., 2008), DCMs of electrophysiological data (Moran, Pinotsis, & Friston, 2013), or DCMs for layered fMRI signals (Heinzle, Koopmans, den Ouden, Raman, & Stephan, 2016). MCMC is also useful when extending DCM to more complex hierarchical models (Raman, Deserno, Schlagenhauf, & Stephan, 2016), in which the derivation of update equations for VB becomes difficult (but see Yao et al., 2018). MCMC is asymptotically exact and only assumes that the posterior distribution can be evaluated up to a multiplicative constant. However, in practice, its computational cost often leads to prohibitively long computation times for the datasets and models commonly encountered in neuroimaging. Furthermore, in contrast to VB, MCMC does not provide an estimate of the model evidence by default.

While several MCMC-based strategies for computing the LME in neuroimaging applications have been explored (e.g., Aponte et al., 2016; Penny & Sengupta, 2016; Raman et al., 2016), one particularly powerful and theoretically attractive MCMC variant is thermodynamic integration (TI). This method, like VB, rests on the concept of free energy and has been proposed as gold standard for LME estimation (Calderhead & Girolami, 2009; Lartillot & Philippe, 2006). Despite strong theoretical advantages, so far, the computational costs of TI have prevented its practical use in neuroimaging.

This paper introduces the reader to thermodynamic integration (TI) and its application to DCM. In contrast to existing tutorials on TI (Annis, Evans, Miller, & Palmeri, 2019), we provide an in-depth discussion of the theoretical foundations of TI and relate the tutorial specifically to DCM as a generative model that is frequently used in contemporary neuroimaging analyses. Our discussion covers the key concept of free energy starting from its historical origin in statistical physics, with the aim of conveying a deeper understanding of this method that goes beyond a purely technical treatment. In the second part, we present a series of examples involving both synthetic and real-world datasets. These include (1) a validation dataset based on a linear regression model with analytically tractable LME used to verify the accuracy of TI, (2) a synthetic fMRI dataset where the true data-generating model is known for each observation, and (3) a real-world fMRI dataset used to demonstrate LME estimation for nonlinear DCM.

In addition to showcasing the application of TI, these examples also serve to demonstrate that, given an efficient implementation and hardware capable of parallel processing, the challenge of high computational demand of TI can be overcome successfully. The software implementation of TI and DCM used in this paper is available as part of the open-source toolbox TAPAS (Translational Neuromodeling Unit, 2014).

To keep this paper short and yet accessible to a broad audience, summaries of key topics such as DCM, Bayesian model selection (BMS), or Markov chain Monte Carlo (MCMC) are offered in the supplementary material (see sections S1, S2 and S3).

## Thermodynamic Integration and the origin of free energy

This section introduces TI from a statistical physics perspective. Statistical physics is a branch of physics that uses methods from probability theory and statistics to characterize the behavior of physical systems. One of the key concepts in statistical physics is that the probability of a particle being in a given state follows a probability density, and that all physically relevant quantities can be derived once this distribution is known. Starting from the free energy, we show how key concepts from information theory have developed from their counterparts in statistical physics, motivating the use of TI and providing a link to the variational Bayes approach conventionally used in DCM to approximate the log model evidence (LME).

### Free energy: A perspective from statistical physics

In thermodynamics, the analogue of the model evidence is the so-called partition function *Z* of a system that consists of an ensemble of particles in thermal equilibrium. A classical discussion of the relationships presented here can be found in Jaynes (1957) and a more modern perspective in Ortega and Braun (2013). For example, let us consider an ideal monoatomic gas, in which the kinetic energy

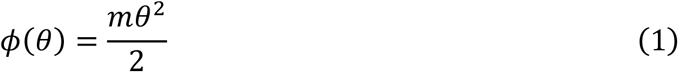

of individual particles is a function of their velocity *θ* and mass *m*. If the system is large enough, the velocity of a single particle can be treated as a continuous random variable. The internal energy *U* of this ideal gas is proportional to the expected energy per particle. It is computed as the weighted sum of the energies *ϕ*(*θ*) associated with all possible velocities, where the weights are given by the probability *q*(*θ*) of the particle being at a certain velocity:

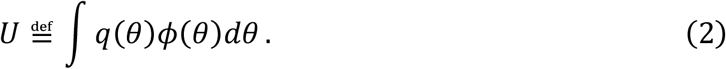

A second important quantity in statistical physics is the entropy *S* of *q*:

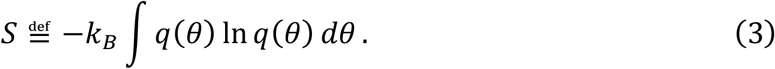

Here, *K*_*B*_ is the Boltzmann constant with units of energy per degree temperature. For an isolated system (i.e., no exchange of matter or energy with the environment), the second law of thermodynamics states that its entropy can only increase or stay constant. Thus, the system is at equilibrium when the associated entropy is maximized, subject to the constraint that the system’s internal energy is constant and equal to *U*, and that *q* is a proper density, i.e.: *q*(*θ*) ≥ 0 and *∫q*(*θ*)*dθ = 1*.

This constrained maximization problem can be solved using Lagrange multipliers (for the derivation see the supplementary material S4), yielding the following distribution:

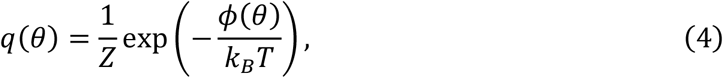

where *Z* is referred to as the partition function of the system:

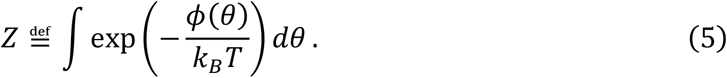

In a closed system, the Helmholtz free energy *F*_*H*_ is defined as the difference between the internal energy *U* of the system and its entropy *S* times the temperature *T*:

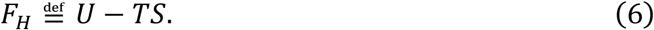

The Helmholtz free energy corresponds to the work (i.e., non-thermal energy in joules that is passed from the system to its environment) that can be attained from a closed system. From Eq. 6, we see that the system with constant internal energy *U* is at equilibrium (i.e., maximum entropy) when the Helmholtz free energy is minimal. Substituting the internal energy (Eq. 2), the entropy (Eq. 3) and the expression of *q* (Eq. 4) into Eq. 6, it follows that the log of the partition function corresponds to the negative Helmholtz free energy divided by *K*_*B*_*T*:

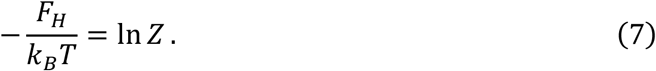

### Free energy: A perspective from statistics

In order to link perspectives on free energy from statistical physics and (Bayesian) statistics, we assume that the system is examined at a constant temperature *T* such that the term *K*_*B*_*T* equals unity (normalization of temperature), allowing us to move from a physical perspective on free energy (expressed in joules) to a statistical formulation (expressed in information units proportional to bits). This is the common convention in the statistical literature, and thereby, all quantities become unit-less information theoretic terms. Under this convention, the physical concept of free energy described above gives rise to an analogous concept of free energy in statistics when the energy function is given by the negative log joint probability *-ln p*(*y, θ*|*mm*) (Neal & Hinton, 1998):

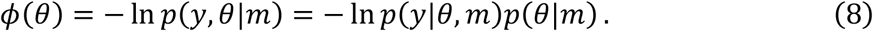

Hence, the log joint (which fully characterizes the system) takes the role of the kinetic energy in the ideal gas example above.

Inserting the expression for *ϕ* (Eq. 8) into Eq. 4, reveals that the equilibrium distribution of the system is the posterior distribution (i.e., the normalized joint probability):

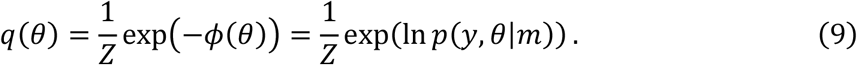

Based on this result, we can define the information theoretic version of the Helmholtz free energy as:

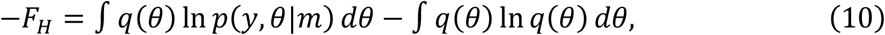

In analogy to Eq. 6, the first term on the right hand side is an expectation over an energy function (cf. Eq. 8); while the second term represents the Shannon entropy *S*_*Shannon*_ = −*∫q*(*θ*) *ln q*(*θ*) *dθ*. Notably, under the choice of the energy function in Eq. 8, the partition function (Eq. 5) corresponds to the normalization constant of the joint probability *p*(*y, θ*|*m*). Comparing with Eq. 7, we see that the negative free energy is equal to the log model evidence (LME):

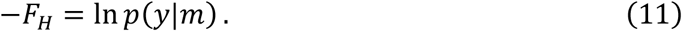

Replacing the joint in Eq. 10 by the product of likelihood and prior, the negative free energy can be decomposed into two terms that have important implications for evaluating the goodness of a model:

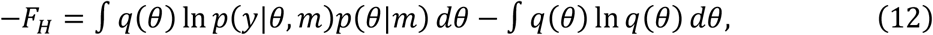

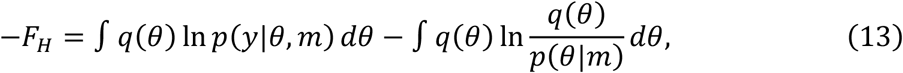

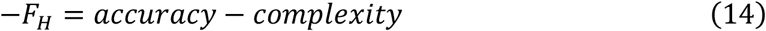

The first term (the expected log likelihood under the posterior) represents a measure of model fit or accuracy. The second term corresponds to the Kullback-Leibler (KL) divergence between the posterior and the prior, and can be viewed as an index of model complexity. Hence, maximizing the negative free energy (log evidence) of a model corresponds to finding a balance between accuracy and complexity. We will turn to this issue in more detail below and examine variations of this perspective under TI and VB, respectively.

In the following, we will explicitly display the sign of the negative free energy for notational consistency. In order to highlight similarities with statistical physics and the concepts of energy and potential, we will continue to express the free energy as a functional of a (possibly non-normalized) log density, such that

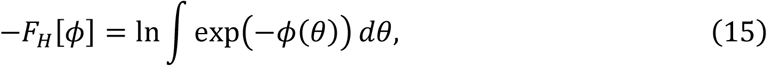

where *ϕ*(*θ*) is equivalent to an energy or potential depending on *θ*. Fig. 1 summarizes the conceptual analogies of free energy between statistical physics and Bayesian statistics.

**Figure 1:**
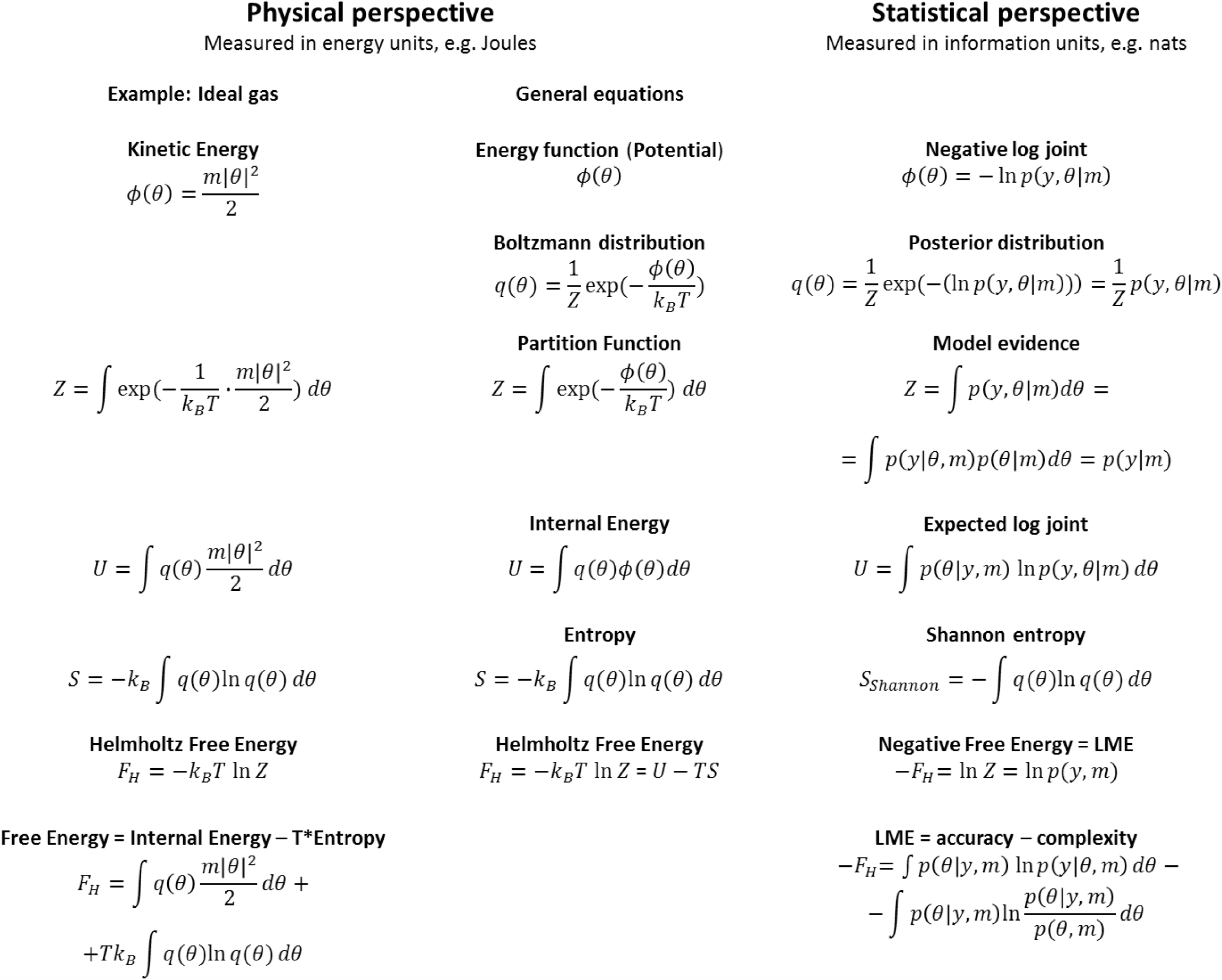
Analogies between concepts of free energy in statistical physics and Bayesian statistics.

### Thermodynamic integration (TI)

We now turn to the problem of computing the negative free energy. As is apparent from Eq. 15, the free energy contains an integral over all possible *θ*, which is usually prohibitively expensive to compute and thus precludes direct evaluation. The basic idea behind TI is to move in small steps along a path from an initial state with known *F*_*H*_ to the equilibrium state and add up changes in free energy for all steps (Gelman & Meng, 1998). This idea was initially introduced in statistical physics to compute the difference in Helmholtz free energy between two states of a physical system (Kirkwood, 1935). Other examples for the application of TI in statistical physics are presented in Landau (2015).

In Bayesian statistics, the same idea can be used to compute the LME of a model *m*. This is because the difference in free energy associated with two potentials corresponding to the negative log prior *ϕ*_0_(*θ*) = −*ln p*(*θ*|*m*) and the negative log joint *ϕ*(*θ*) = −*ln p*(*y*|*θ, m*) −*ln p*(*θθ*|*mm*) (cp. Eq. 8) equals the LME. More precisely, provided the prior is properly normalized, i.e., *∫p*(*θ*|*m*)*dθ =*1, substituting *ϕ*_0_ and *ϕ* into Eq. 15 yields

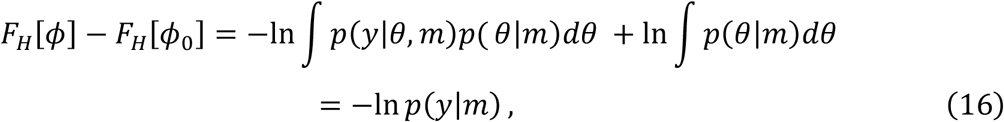

The goal is now to construct a piecewise differentiable path connecting prior and posterior and then compute the LME by integrating infinitesimal changes in the free energy along this path. A smooth transition between *F*[*ϕ*] and *F*[*ϕ*_0_] can be constructed by the power posteriors *p*_*β*_(*θ*|*y, m*) (see Eq. 18 below) which are defined by the path *ϕ*_*β*_:

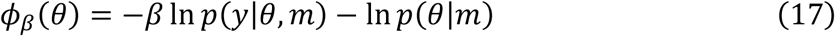

with *β ϵ* [0,1], such that *ϕ*_1_ =*ϕ*. In the statistics literature, *β* is usually referred to as an inverse temperature because it has analogous properties to physical temperature in many aspects. We will use this terminology and comment on the analogy in more detail below.

The power posterior is obtained by normalizing the exponential of −*ϕ*_*β*_(*θ*):

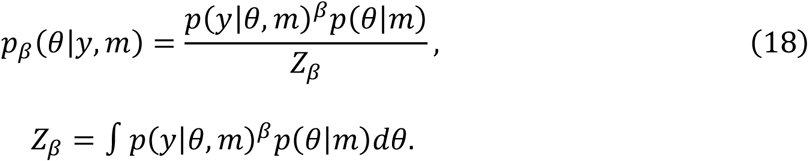

Combining this definition with Eq. 16, the LME can be written in terms of an integral over an expectation with respect to the power posterior

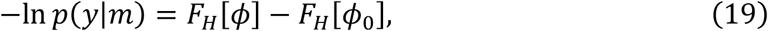

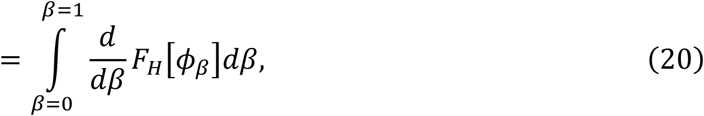

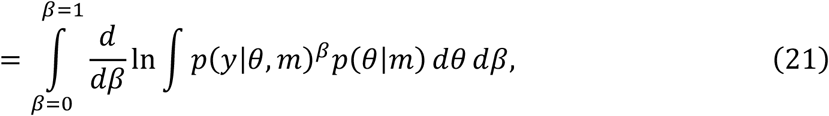

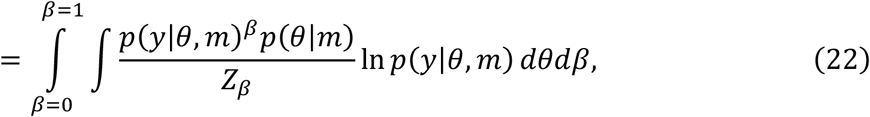

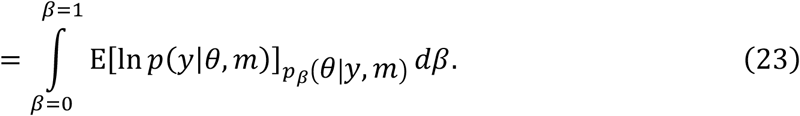

which we refer to as the basic or fundamental TI equation (Gelman & Meng, 1998).

Notably, the TI equation can also be understood in terms of the definition of the free energy (Eq. 13) by noting that the latter can be written as the sum of an expected log likelihood and a cross-entropy term (KL divergence between power posterior and prior):

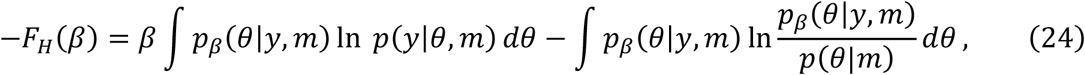

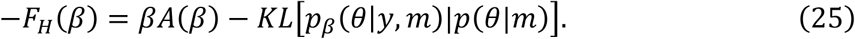

The first term, *A*(*β*) = −*∂F*_*H*_*/∂β*, is referred to as the accuracy of the model (see, for example, Penny, Stephan, Mechelli, & Friston, 2004a; Stephan, Penny, Daunizeau, Moran, & Friston, 2009), while the second term constitutes a complexity term. Note that Eq. 25 is typically presented in the statistical literature for the case of *β = 1* and describes the same accuracy vs. complexity trade-off previously expressed by Eq. 13, but now from the specific perspective of TI.

Fig. 2 shows a graphical representation of the relation conveyed by the fundamental TI equation (Eq. 23) and Eq. 25. For any given *β*, the negative free energy at this position of the path −*F*_*H*_(*β*) can be interpreted as the signed area below the curve *Aβ*(*β*) = −*∂F*_*H*_*/∂β* (i.e., the integral over *A*(*⋅*) from 0 to *β*), whereas the term *β* × *A*(*β*) is the rectangular area below *ββ*(*ββ*). Eq. 25 shows that the area *βA*(*β*) *+ F*_*H*_(*β*) is the KL divergence between the corresponding power posterior and prior.

**Figure 2:**
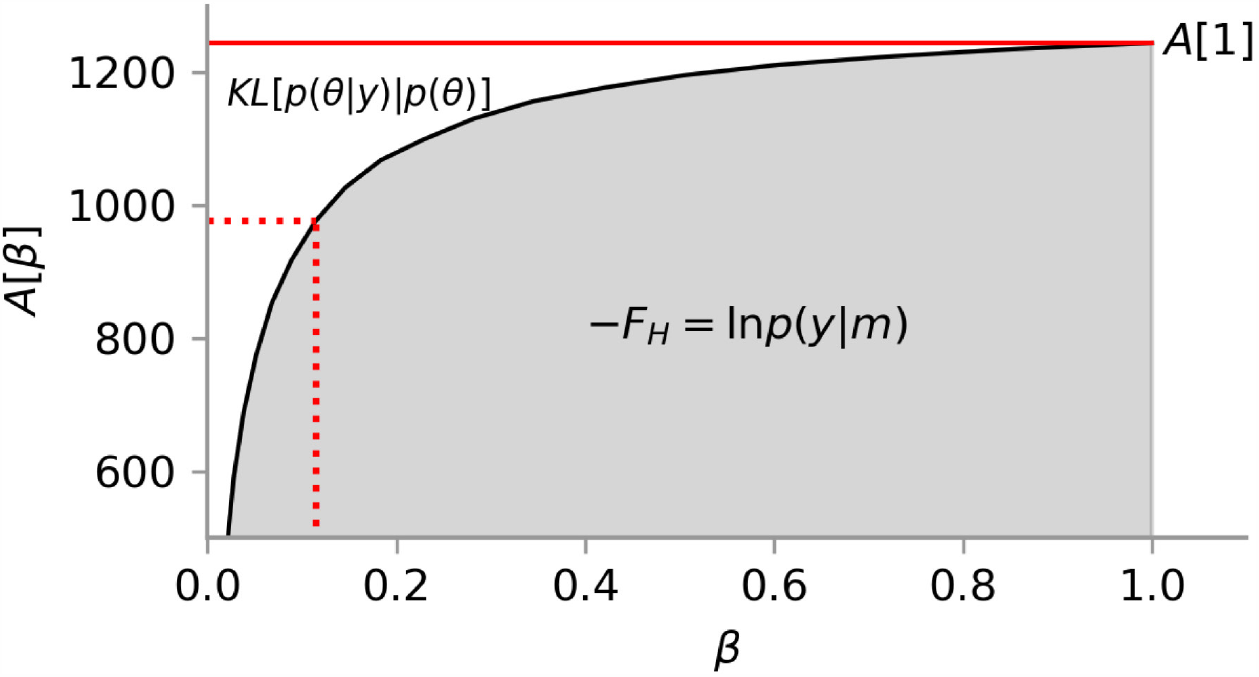
Graphical representation of the TI equation. The free energy is equal to the *signed* area below *A=* −*∂F*_*H*_ */∂β*, and thus the area *A*(1) *+ F*_*H*_ is equal to the KL divergence between posterior and prior. The same relation holds for any *β* ∈ [0,1].

This relationship holds because, for the power posteriors (Eq. 18), *A*(*ββ*) is a monotonically increasing function of *ββ*. This is due to the fact that

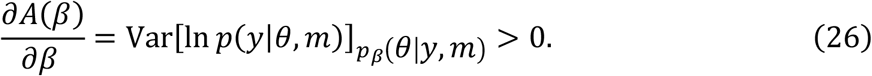

See Lartillot and Philippe (2006) for a derivation of this property. From this it follows that the negative free energy is a concave function along *β*.

The theoretical considerations highlighted above and the relation to principles of statistical physics render TI an appealing choice for estimating the LME. However, the question remains how the LME estimator in Eq. 23 can be evaluated in practice. To solve this problem, TI relies on Monte Carlo estimates of the expected value *E*[*ln p*(*y*|*θ, m*)]*p*_*β*_(*θ*|*y,m*) in Eq. 23:

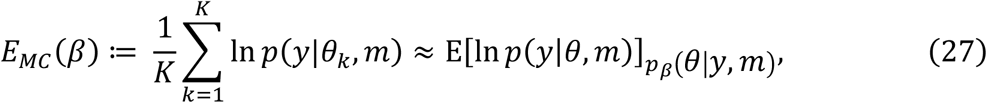

where samples *θ*_*K*_ are drawn from the power posterior *p*_*β*_(*θ*|*y, m*). The remaining integral over *β* in Eq. 23 is a one dimensional integral, which can be computed through a quadrature rule using a predefined temperature schedule for *β* (0 = *β*_0_ < *β*_1_ < … < *β*_*N*−1_ < *β*_*N*_ =*1*):

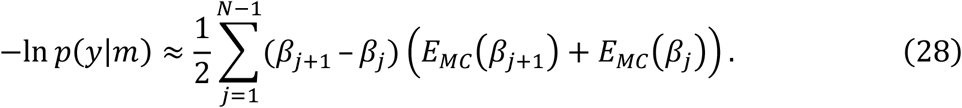

The optimal temperature schedule in terms of minimal variance of the estimator and minimal error introduced by this discretization was outlined previously in the context of linear models by Gelman and Meng (1998) and Calderhead and Girolami (2009).

Note that each step *β*_*j*_ in the temperature schedule requires a new set of samples *θ*_*K*_ to be drawn from the respective power posterior *pβ*_*j*_ (*θ*|*y, m*), contributing to the high computational complexity of TI. However, since these sets of samples are independent from each other, this can in principle be done in parallel, provided suitable soft- and hardware capabilities are available. An efficient way to realize such a parallel sampling procedure is to adopt a population MCMC approach in which MCMC sampling is used to generate, for each *β*_*j*_, a chain of samples from the respective power posterior *pβ*_*j*_ (*θ*|*y, m*). In addition, chains from neighboring *β*_*j*_ in the temperature schedule are allowed to interact by means of a “swap” accept-reject (AR) step (Swendsen & Wang, 1986). This increases the sampling efficiency and speeds up convergence of the individual MCMC samplers. For readers unfamiliar with Monte Carlo methods, a primer on (population) MCMC is provided in the supplementary material S3. For a detailed treatment, we refer to McDowell, Dyckman, Austin, and Clementz (2008) and Calderhead and Girolami (2009).

So far, the computational requirement of sampling from an ensemble of distributions (one for each value of *β*) has limited the application of TI to high performance computing environments and prevented its widespread use in neuroimaging. Luckily, the increase in computing power of stand-alone workstations and the proliferation of graphical processing units (GPU), coupled with efficient population MCMC samplers, offer possibilities to overcome this bottleneck, which will be demonstrated below for a selection of three examples involving synthetic and real-world datasets. First, however, we will complete the theoretical overview by briefly explaining the formal relationship between TI and variational Bayes.

### Variational Bayes

Variational Bayes (VB) is a general approach to approximate intractable integrals with tractable optimization problems. Importantly, this optimization method simultaneously yields an approximation to the posterior density and a lower bound to the LME.

The fundamental equality which underlies VB is based on introducing a tractable density *q*(*θ*) to approximate the posterior *p*(*θ*|*y, m*).

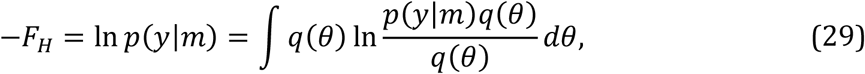

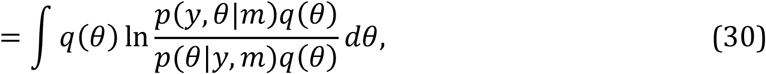

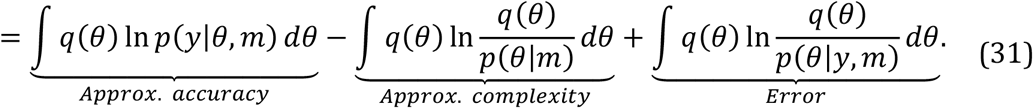

The last term in Eq. 31 is the KL divergence between the approximate density *q* and the unknown posterior density; this encodes the error or inaccuracy of the approximation. Given that the KL divergence is never negative, the first two terms in Eq. 31 represent a lower bound on the log evidence −*F*_*H*_, and in the following we will refer to it as the negative variational free energy −*F*_*VB*_.

In summary, the relation between the information theoretic version of Helmholtz free energy −*F*_*H*_, log model evidence *ln pp*(*yy*|*mm*), and variational negative free energy −*F*_*VB*_is therefore

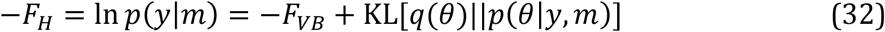

We highlight this relationship because many readers are rightfully confused that the term ‘negative free energy’ is sometimes used in the literature to denote the logarithm of the partition function *Z* itself (i.e., −*F*_*H*_), as we have done above, and sometimes to refer to a lower bound approximation of it (i.e., −*F*_*VB*_). This is because the variational free energy −*F*_*VB*_ becomes identical to the negative free energy −*F*_*H*_ when the approximate density *q* equals the posterior and hence their KL divergence becomes zero. In this special case

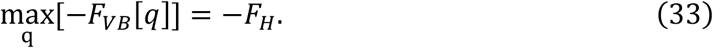

To maintain consistency in the notation, we will distinguish −*F*_*H*_ and −*F*_*VB*_ throughout the paper. VB aims to reduce the KL divergence between *q* and the posterior density by maximizing the lower bound −*F*_*VB*_ as a functional of *q*:

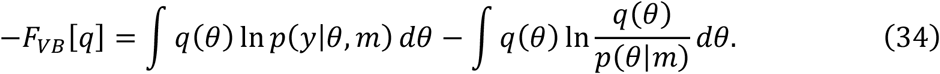

When the functional form of *q* is fixed and parametrized by a vector *η*, VB can be reformulated as an optimization method in which *η* is updated according to gradient *∂F*_*VB*_[*q*(*θ*|*η*)]*/∂η* (Friston et al., 2007). Thus, the path followed by *η* during optimization can be formulated as

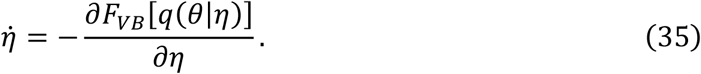

This establishes a connection between TI and VB. In the former, the path of *η* corresponds to the path of *β* from 0 to 1, which was selected a priori with the conditions that

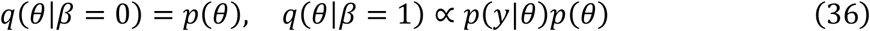

and the gradients

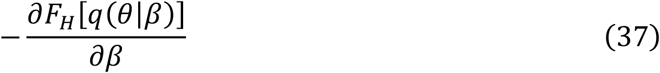

are used to numerically compute the free energy.

Different VB algorithms are defined by the particular functional form used for the approximate posterior. In the case of DCM, it is so far most common to use Variational Bayes under the Laplace approximation (VBL). A summary of VBL for DCM is available in the supplementary material S5, while an in-depth treatment is provided in Friston et al. (2007).

## Evaluating the accuracy of TI

In this section, we investigate the accuracy of LME estimates obtained with TI, and compare the performance of TI to that of two other sampling-based LME estimators, the prior arithmetic mean (AME) and posterior harmonic mean (HME) estimators. In contrast to TI, which requires sampling from an ensemble of distributions (see Eq. 28), AME and HME only require samples from the prior or posterior distributions, respectively. Hence, these two methods, which are described in detail in the supplementary material S6, are computationally significantly less demanding than TI.

For the purpose of this comparison, we turn to a Bayesian linear regression model with normal prior and likelihood. This is a useful case for benchmarking because the LME can be computed analytically. This model is described by the following prior and likelihood function:

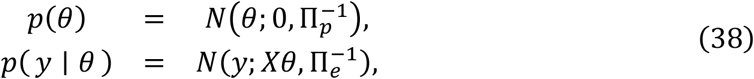

where *θ* is the [*p* × 1] vector of regression coefficients, *y* is the [*M* × 1] vector of data points, *X* is the [*M*× *p*] design matrix, and 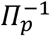 and 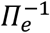 are the covariance matrices of the

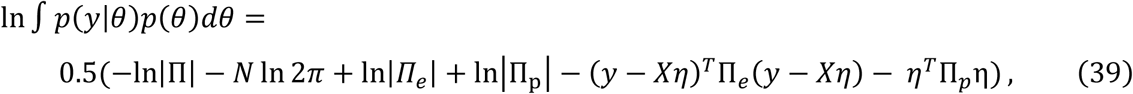

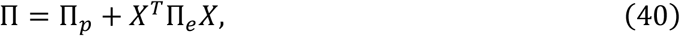

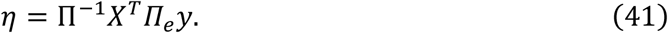

For our simulations, we chose *M = 100*, 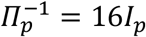 and 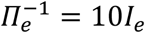, where *I*_*p*_ and *I*_*e*_ are the corresponding identity matrices. The design matrix was chosen to have a block structure equivalent to a design for a one-way ANOVA with *p* levels (for those values of *p* that do not exactly divide by *M*, the excess data points were assigned to the last cell). Synthetic data where generated by sampling from the generative model defined in Eq. 38.

By varying *p* from 2 to 32 in steps of 2, we created a series of models with increasing dimensionality. For each value of *pp*, we repeated the data generation process 10 times, drawing a new set of values for the regression parameters *θθ* each time from the prior, and generating observations *y* according to the likelihood. We then estimated the LME using TI, AME and HME, and compared the estimates against the analytically computed LME.

The TI approximation to the LME was computed using 64 chains with a 5th order annealing schedule, i.e. a temperature schedule with 64 steps *β*_*j*_, with step size chosen according to a fifth order power rule (Calderhead & Girolami, 2009). In each chain, we generated 6000 samples. We then computed AME based on the samples from the prior density and HME based on samples from the posterior. Fig. 3 shows the error in the LME estimates as a function of the number of model parameters for the three approaches. Consistent with previous reports, the results show that HME overestimated the LME, while AME underestimated it (Lartillot & Philippe, 2006). Only TI provided good estimates over the full range of models. This indicates that, despite comparing unfavorably in terms of computational efficiency, TI should still be preferred in practice due to the large estimation error of AME and HME, especially for higher-dimensional models (but see Penny and Sengupta (2016) for variants of AME and HME that solve some of the issues).

**Figure 3:**
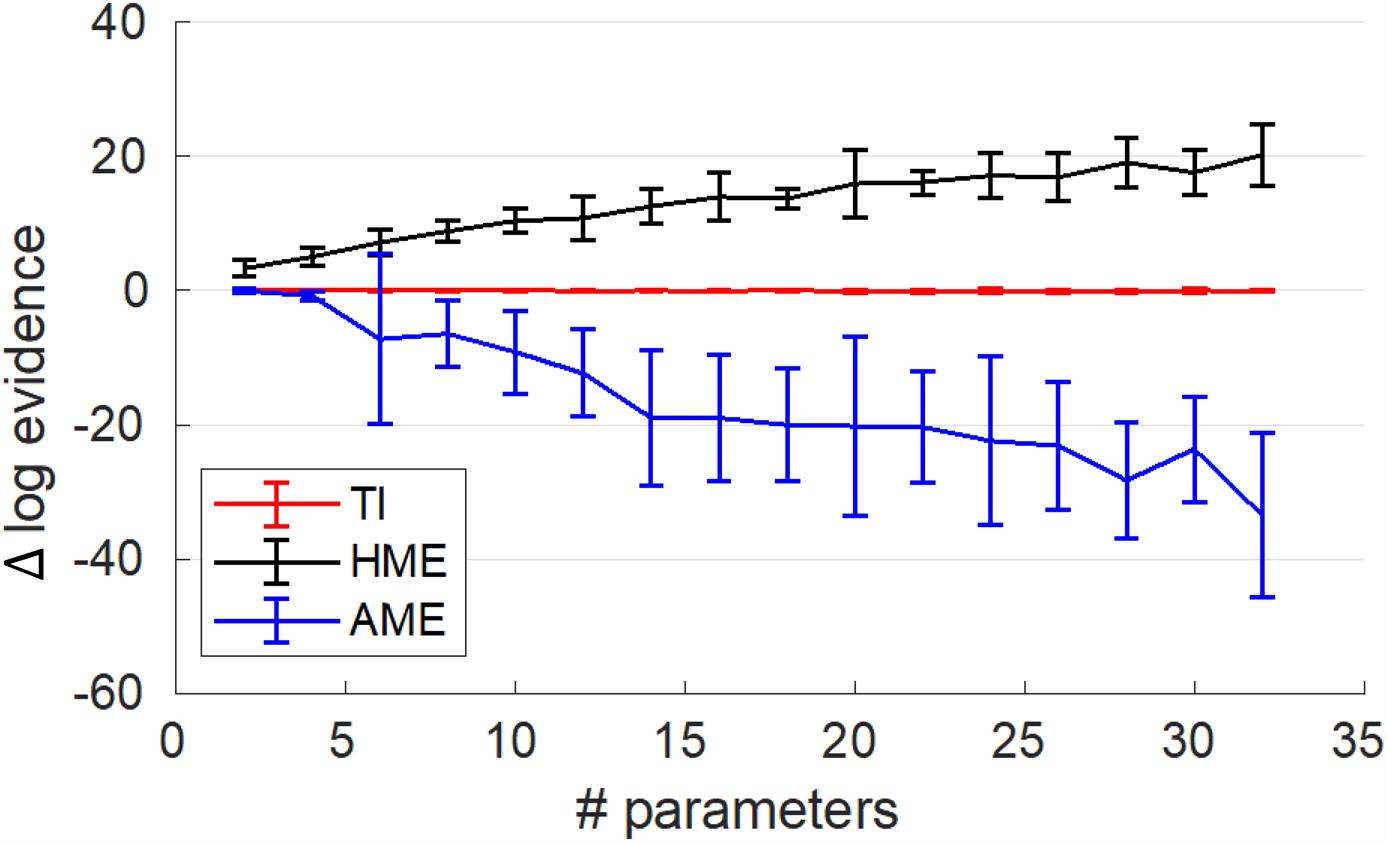
Error in estimating the log evidence of linear models for three different sampling approaches. The curves show mean and standard deviation (error bars) over ten runs at each value of *p* (number of GLM parameters) for thermodynamic integration (TI), posterior harmonic mean estimator (HME) and prior arithmetic mean estimator (AME).

## Application to DCM

Having established the accuracy of TI as an estimator for the LME in a case where the analytic solution for the LME is available, we now turn to the case of LME estimation and model selection in the context of DCM. For this purpose, we discuss two example applications. The first example considers a simulated dataset where the true model that generated each observation is known. This serves to determine the ability of TI to identify the data-generating model. In the second example, we analyze the “attention to motion” fMRI dataset (Buchel, 1997), which has been analyzed by numerous previous methodological studies. Primers on DCM and Bayesian model selection are provided in the supplementary material.

### DCM: Simulated data

In the first experiment, we used simulated data from 5 DCMs (linear: model 1; bilinear: models 2 to 4; nonlinear: model 5) with two inputs (*u*_1_ and *u*_2_). The DCMs are displayed in Fig. 4 and are available for download via the ETH Research Collection (ETH Zurich, 2020). The numerical values of the connectivity matrices are listed in the supplementary material S7. The BOLD signal data were simulated assuming a repetition time (TR) = 2s and 720 scans per simulation. The driving inputs were entered with a sampling rate of 2.0*Hz*. Simulated time series were corrupted with Gaussian noise yielding a signal-to-noise ratio (SNR) of 1.0. Here, SNR was defined as the ratio of signal standard deviation to noise standard deviation (Welvaert & Rosseel, 2013). This means that our simulated data contained identical amounts of noise and signal, representing a relatively challenging SNR scenario.

**Figure 4:**
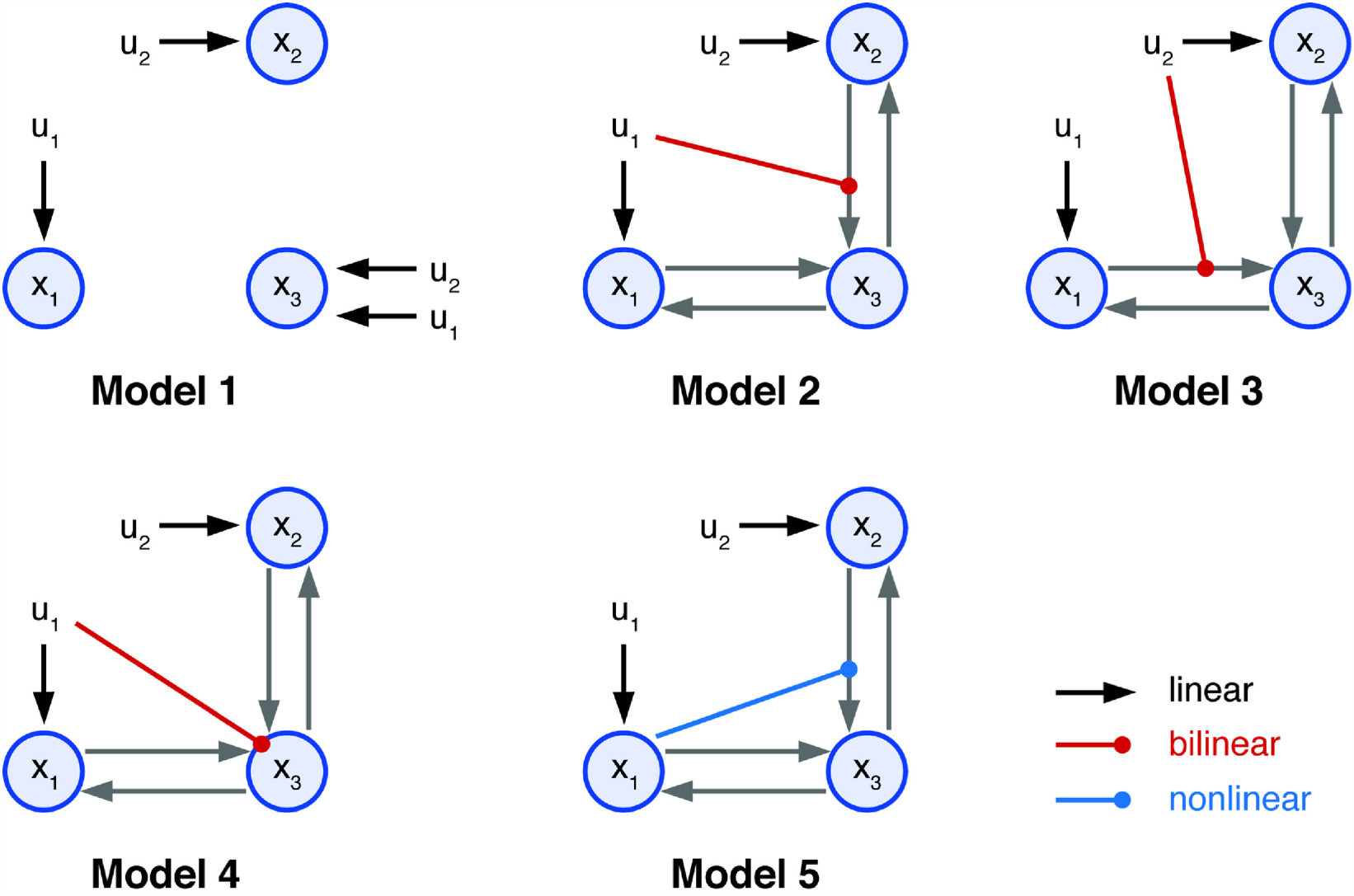
Illustration of the five simulated 3-region DCMs used for cross-model comparison. Self-connections are not displayed. The variables u_1_ and u_2_ represent two different experimental conditions or inputs. All models represented different hypotheses of how the neuronal dynamics in area x_3_ could be explained in terms of the two driving inputs and the effects of the other two regions x_1_ and x_2_. Model m_1_ can be understood as a ‘null hypothesis’ in which the activity of all the areas can be explained by the driving inputs. Models m_2_ and m_3_ correspond to two forms of bilinear effect on the forward connection of areas x_1_ and x_2._ Model m_4_ represents the hypothesis that input u_1_ affects the self-connection of area x_3_ (not displayed). Model m_5_ represents a non-linear interaction between regions x_1_ and x_2_. Endogenous connections are depicted by gray arrows, driving inputs by black arrows, bilinear modulations by red arrows and nonlinear modulations by blue arrows

For each model, we generated 40 different datasets with different instantiations of Gaussian noise, such that the underlying time series remained constant. We then counted how often the data-generating model was assigned the largest model evidence and compared the ensuing values across the different estimators (i.e., AME, HME, TI, VBL). Notably, the absolute value of the log evidence of a given model is irrelevant for model scoring; instead, its difference to the log evidence of other models is decisive.

In a pretesting phase, we found that TI generated stable estimates of the LME using 64 chains. All simulations were executed with a burn-in phase of 1 × 10^4^ samples, followed by an additional 1 × 10^4^ samples used for analysis. We evaluated the convergence of the MCMC algorithm by examining the potential scale reduction factor 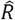 (Gelman & Rubin, 1992) for samples of the log likelihood of all chains. We found that 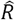 was below 1.1 in all but a few instances, indicating convergence. Estimated LME values are displayed in Fig. 5 and Table 1. Consistent with the linear model analysis in the previous section, the HME was always higher and the AME always lower than the TI estimate of the LME. VBL estimates were close to the TI estimate. To test for significant differences in accuracy of recovering the correct model by the different algorithms, *χ*^2^ tests were employed. TI and VBL were not significantly different (*χ*^2^ = 0.3, *p* = 0.56), but both TI and VBL (not shown) were significantly better than AME (*χ*^2^ = 189.5, *p* < 10^−5^) and HME (*χ*^2^ = 25.4, *p* < 10^−5^).

**Table 1:** LME estimated with TI and VBL. Tables display the LME (summed across 40 simulations) of each combination of inverting and generating models. Columns have been normalized by the lowest LME: according to TI. Columns on the right and left tables share the same normalization and their absolute values can be directly compared. On most, but not all occasions, VBL underestimated the LME compared to TI. However, for both VBL and TI the data-generating model obtained the highest LME (diagonals).

|  |  | TI |  |  |  |  | VBL |  |  |  |  |  |
| --- | --- | --- | --- | --- | --- | --- | --- | --- | --- | --- | --- | --- |
|  |  | Generating model |  |  |  |  | Generating model |  |  |  |  |  |
| | | $m_1$ | $m_2$ | $m_3$ | $m_4$ | $m_5$ | | $m_1$ | $m_2$ | $m_3$ | $m_4$ | $m_5$ |
| Inverting model | $m_1$ | 762 | 0 | 0 | 0 | 0 | $m_1$ | 746 | 18 | 16 | -3 | 11 |
| | $m_2$ | 10 | 11155 | 10676 | 4242 | 4359 | $m_2$ | -34 | 11101 | 10614 | 4182 | 4294 |
| | $m_3$ | 6 | 10360 | 11683 | 2903 | 4349 | $m_3$ | -39 | 10298 | 11627 | 2842 | 4288 |
| | $m_4$ | 0 | 10038 | 9102 | 5163 | 4376 | $m_4$ | -49 | 9984 | 9028 | 5097 | 4310 |
| | $m_5$ | 84 | 9045 | 9190 | 2931 | 4430 | $m_5$ | 44 | 9034 | 9181 | 2877 | 4375 |

**Figure 5:**
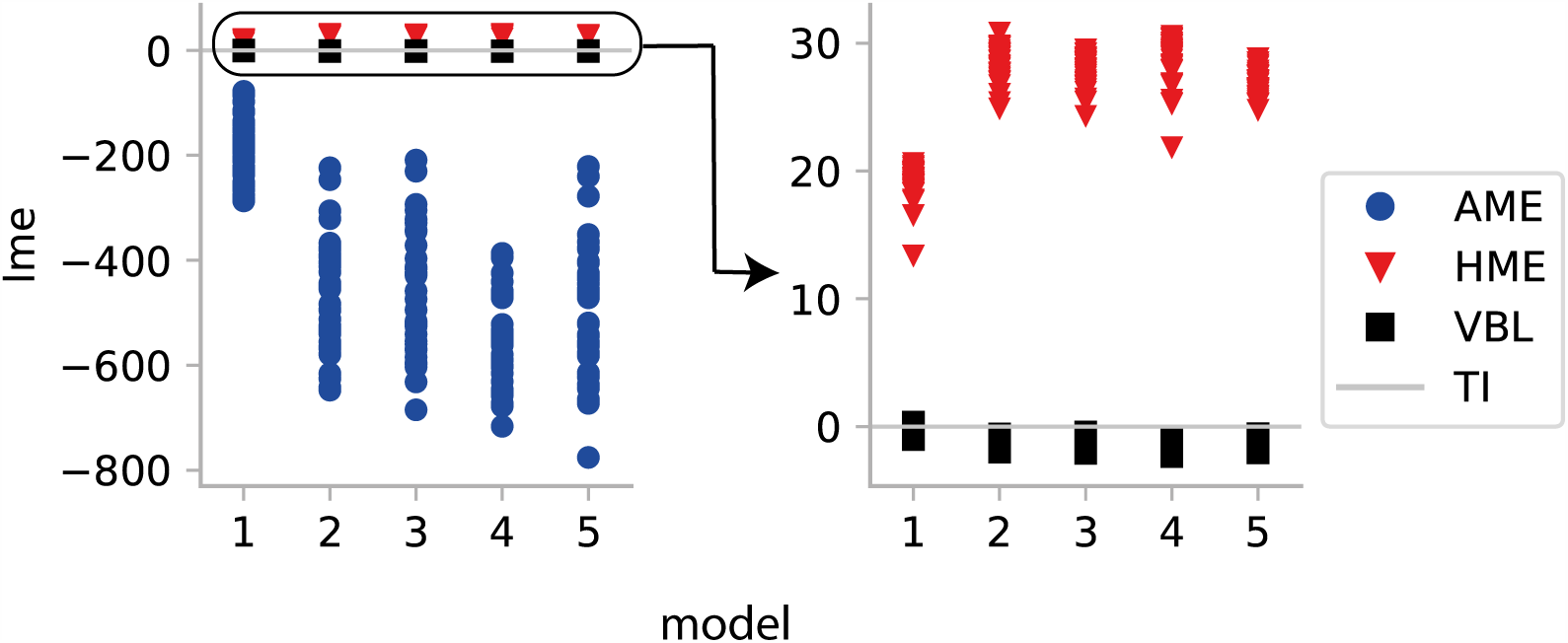
Estimated LME for all models relative to TI when inverted with the corresponding data-generating model under *SNR = 1* for 40 different models. Right panel zooms in the left panel. Red triangles correspond to the HME, blue circles to the AME, and black squares to VBL. HME was always higher and AME always lower than the TI estimate. All LME estimates are shown after subtracting the TI-based estimate for the same model.

We then examined how often the data-generating model was identified correctly by model comparison, i.e., how often it showed the largest LME of all models. Of all estimators, AME failed most frequently to detect the data-generating model (Table 2). HME identified the correct model more consistently (Table 3). Both VBL and TI displayed a similar behavior (Tables 4 and 5), although model *m*_5_ was identified slightly more consistently identified by VBL. However, as displayed in Table 1, according to both inversion schemes, the data-generating model was consistent with the model showing the highest LME.

**Table 2:** Cross-model comparison results for AME in the case of synthetic data (SNR = 1). The row label indicates the data-generating model, the column index is the inferred model.

| AME: Synthetic data |  |  |  |  |  |  |
| --- | --- | --- | --- | --- | --- | --- |
| Inversion | Generation |  |  |  |  |  |
| | | $m_1$ | $m_2$ | $m_3$ | $m_4$ | $m_5$ |
| | $m_1$ | 39 | 14 | 10 | 34 | 29 |
| | $m_2$ | 1 | 13 | 11 | | 3 |
| | $m_3$ | | 6 | 11 | 3 | |
| | $m_4$ | | 3 | 4 | 2 | 5 |
| | $m_5$ | | 4 | 4 | 1 | 3 |

**Table 3:**
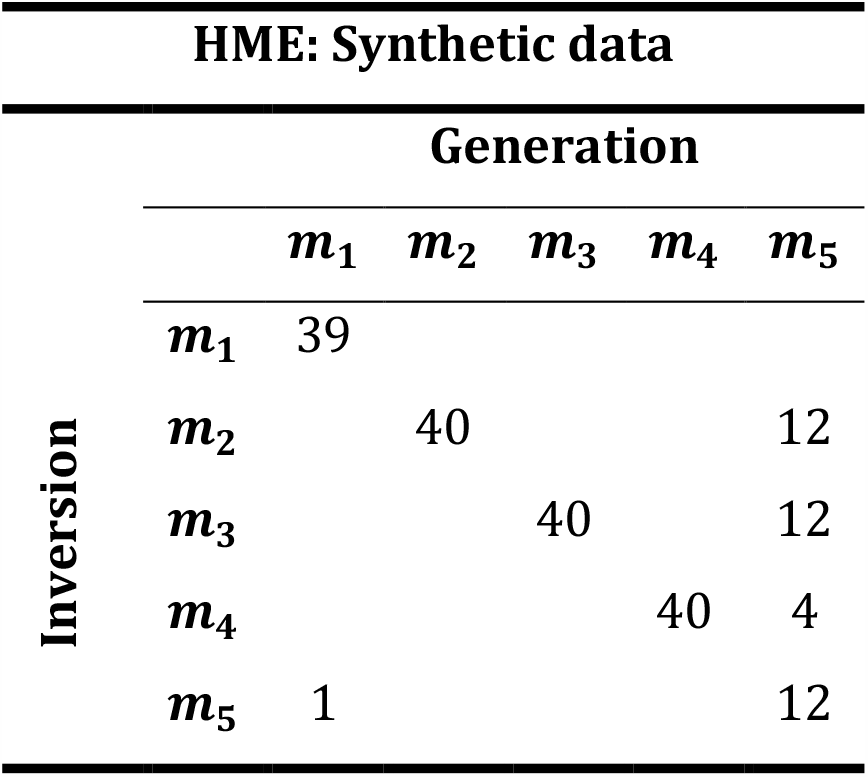
Cross-model comparison results for HME in the case of synthetic data (SNR = 1). The row label indicates the data-generating model, the column index is the inferred model.

**Table 4:**
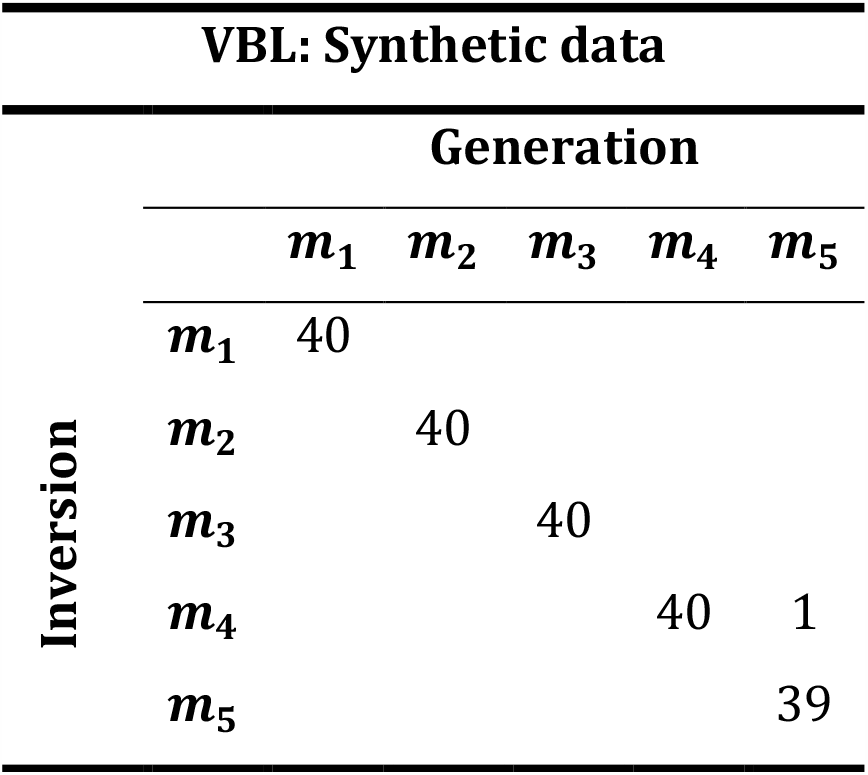
Cross-model comparison results for VBL in the case of synthetic data (SNR = 1). The row label indicates the data-generating model, whereas the column index is the inferred model.

**Table 5:**
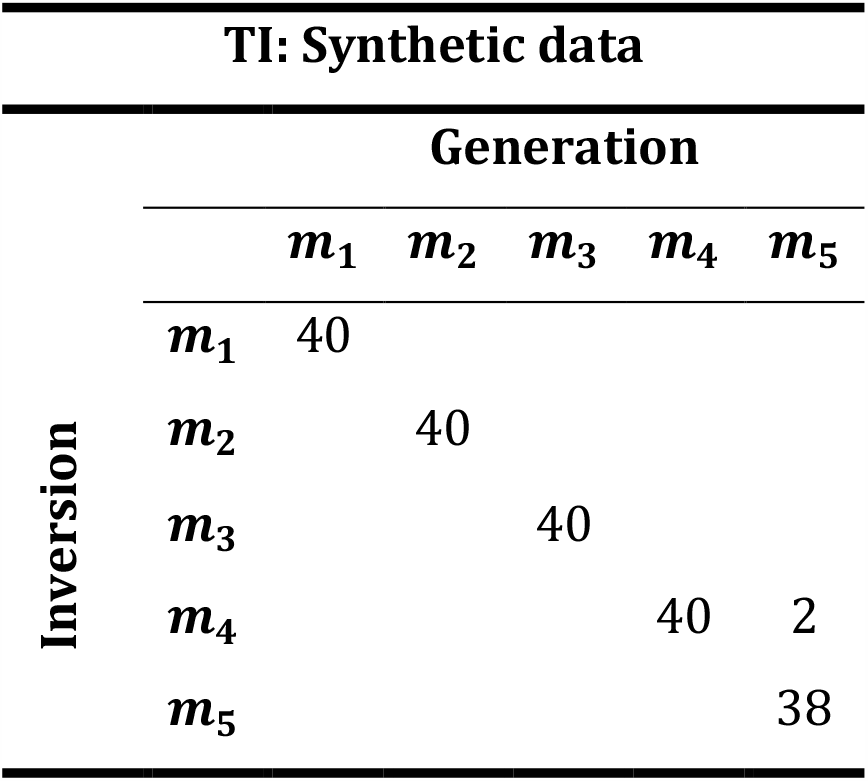
Cross-model comparison results for TI in the case of synthetic data (SNR = 1). The row label indicates the data-generating model, the column index is the inferred model.

### Empirical data: Attention to motion

In this example, we demonstrated TI-based parameter estimation and model comparison for DCM on an empirical dataset. Since the previous two examples have shown that TI consistently outperforms the other sampling-based LME estimators, AME and HME, we limit our comparison to TI and VB, from here on.

For the analysis of empirical data, we selected the “attention to motion” fMRI dataset (Buchel, 1997) that has been analyzed in numerous previous methodological studies (e.g, Friston et al., 2003; Marreiros, Kiebel, & Friston, 2008; Penny et al., 2004a; Penny, Stephan, Mechelli, & Friston, 2004b; Stephan et al., 2008). The original study investigated the effect of attention on motion perception (Buchel, 1997); in particular, the authors examined attentional effects on the connectivity between primary visual cortex (V1), motion-sensitive visual area (V5) and posterior parietal cortex (PPC). In brief, the experimental paradigm consisted of four conditions (all under constant fixation): fixation only (F), presentation of stationary dots (S), passive observation of radially moving dots (N), or attention to the speed of these dots (A). Four sessions were recorded and concatenated, yielding a total of 360 volumes (*T*_*E*_ = 40*ms, TR* = 3.22*s*). Three inputs were constructed using a combination of the three conditions: *stimulus* = *S* + *N* + *A, motion* = *N* + *A, attention* = *A*. Driving inputs were resampled at 0.8*Hz*, requiring a total of 1440 integration steps. Further details of the experimental design and analysis can be found in Buchel (1997).

One reason for selecting this dataset is that Stephan et al. (2008) previously demonstrated that a nonlinear model (model 4 in Fig. 6) had higher evidence than comparable bilinear models (model 1-3 in Fig. 6). This case is of interest for evaluating the quality of different LME estimators, as one would expect that the introduction of nonlinearities represents a challenging case for VBL.

**Figure 6:**
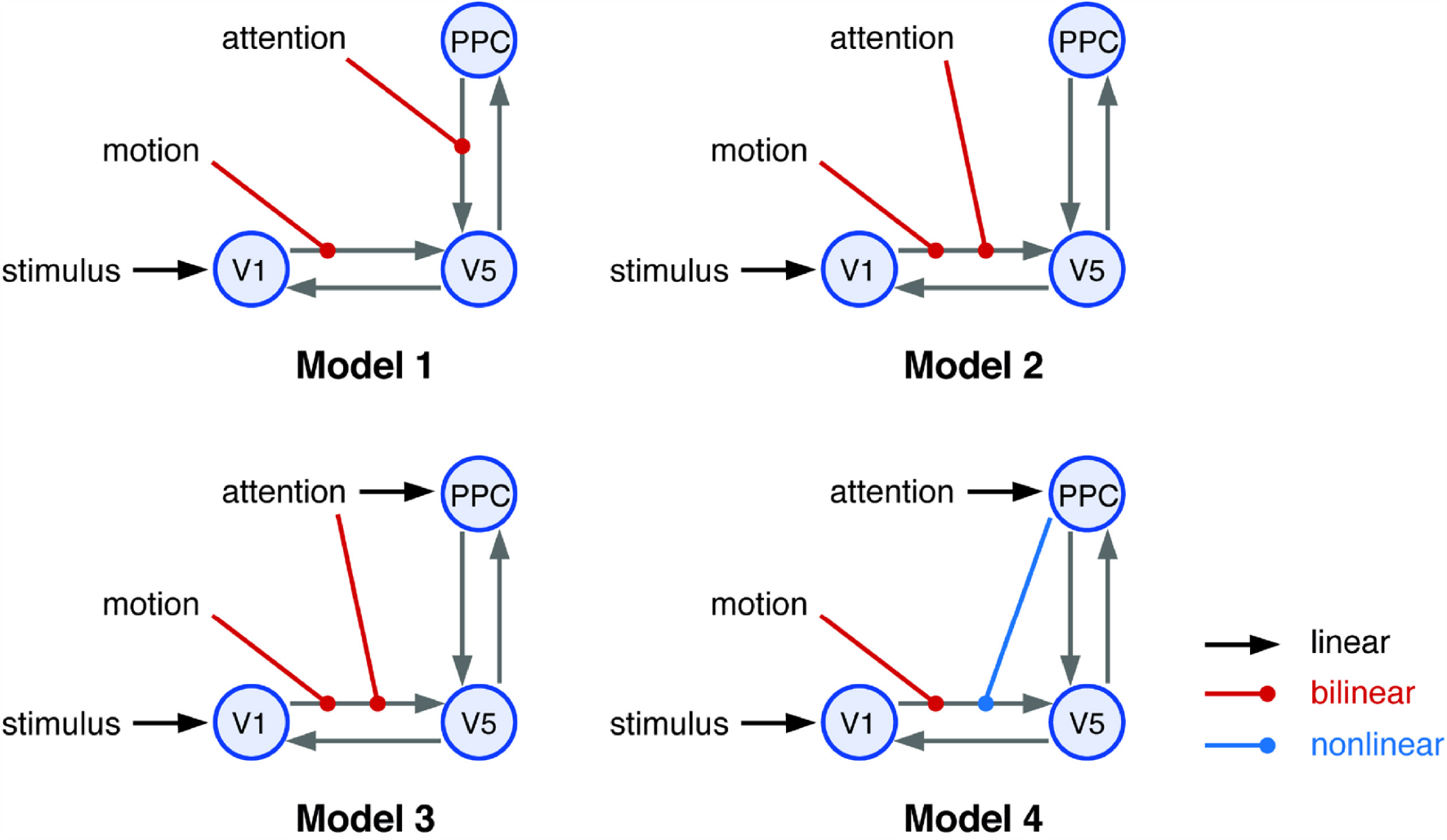
Illustration of the four models used in Stephan et al. (2008) representing different hypotheses of the putative mechanisms underlying attention-related effects in the motion-sensitive area V5. The first three models are bilinear whereas the fourth model is a nonlinear DCM. Endogenous connections are depicted by gray arrows, driving inputs by black arrows, bilinear modulations by red arrows and nonlinear modulations by blue arrows. Inhibitory self-connections are not displayed. V1: primary visual area, V5 = motion sensitive visual area, PPC: posterior parietal cortex.

For TI-based LME estimation, 16 × 10^3^ samples were collected from 64 chains, of which 8 × 10^3^ were discarded in the burn-in phase. The convergence of the algorithm was evaluated using the 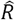 statistic of the samples of the log likelihood of each chain and model. In all but one chain, 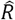 was below 1.1, indicating convergence.

Table 6 summarizes the evidence estimates obtained with TI and VBL. In comparison to previous results see Table 8 in Stephan et al. (2008), three findings are worth highlighting. First, as shown in Table 6, the VBL algorithm reproduced the ranking of models reported in Stephan et al. (2008), although an earlier version of the VBL algorithm with different prior parameters and a different integration scheme was used by Stephan et al. (2008). Moreover, our TI implementation produced the same ranking as the one obtained under VBL.

**Table 6:** Results of model comparison, in terms of log evidence differences with respect to the worst model (m_1_), from Stephan et al. (2008), who used a different prior and integrator as in here.

| $m_1$ | $m_2$ | $m_3$ | $m_4$ |
| --- | --- | --- | --- |
| **Stephan et al. 2008. (VBL)** |  |  |  |
| 0.0 | 3.1 | 5.6 | 13.6 |
| **VBL** |  |  |  |
| 0.0 | 11.4 | 13.4 | 15.2 |
| **TI** |  |  |  |
| 0.0 | 11.5 | 14.8 | 43.5 |

Second, the difference between the VBL free energy estimates and the TI estimates varied considerably across models. To investigate this variability, we compared TI and VBL with regard to the accuracy term. The results are summarized in the lower section of Table 7. Table 6 shows that the discrepancies between VBL and TI varied across models, and the difference was particularly pronounced for the nonlinear model *m*_4_ (>40 log units).

**Table 7:** Log model evidence, accuracy and log likelihood at the MAP estimate using both TI and VBL.

| **Attention to motion dataset** |  |  |  |  |
| --- | --- | --- | --- | --- |
| **Log model evidence** |  |  |  |  |
| | $m_1$ | $m_2$ | $m_3$ | $m_4$ |
| VBL | -1790.0 | -1778.6 | -1776.6 | -1774.8 |
| TI | -1772.6 | -1761.1 | -1757.8 | -1729.1 |
| **Accuracy** |  |  |  |  |
| | $m_1$ | $m_2$ | $m_3$ | $m_4$ |
| VBL | -1547.6 | -1538.5 | -1531.6 | -1530.7 |
| TI | -1525.6 | -1520.2 | -1511.8 | -1483.5 |

Third, while VBL detected the most plausible model, the findings from this dataset suggest that VBL-based inversion of DCMs might not always be fully robust. In particular, the difference between the algorithms could be attributed to the VBL algorithm converging to a local extremum. To assess the differences between TI and VBL more systematically, we initialized each algorithm 10 times from different starting values that were randomly sampled from the prior density. Fig. 7 depicts the estimated model evidence and accuracy and Fig. S1 in the supplementary material S10 displays the predicted BOLD signal. VBL estimates of the accuracy and LME displayed much larger variance than the TI estimates. This suggests that the greater variance of the VBL estimates is due to the propensity of the gradient ascent used in VBL to converge to local maxima.

**Figure 7:**
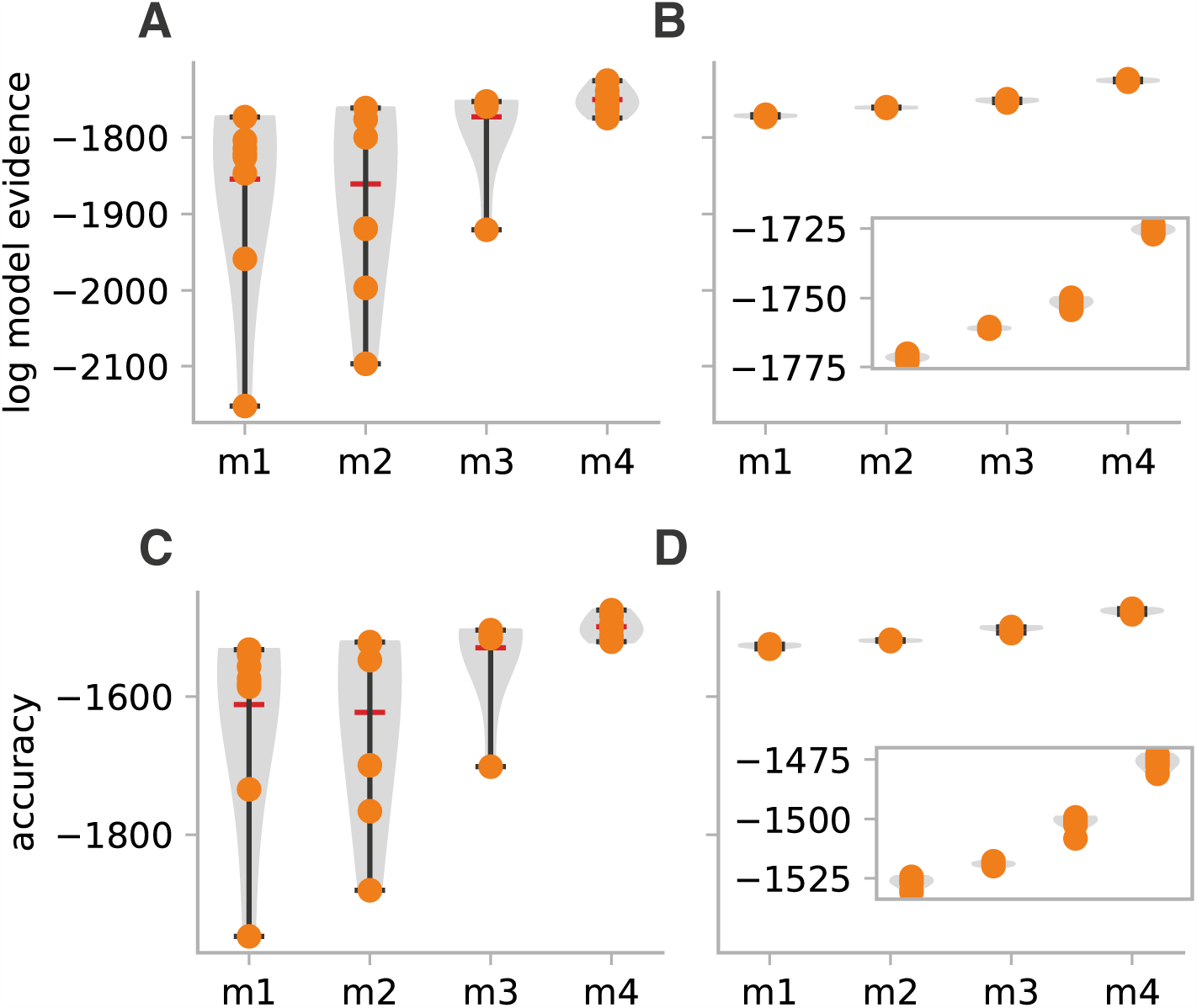
Estimates of the LME and accuracy in the attention to motion dataset after initializing VBL and TI from 10 different starting points (yellow points) drawn from the prior. The inset on the right panel zooms into the range of TI estimates. **A**. LME estimates from VBL. **B**. LME estimates from TI. **C**. Accuracy component of the LME estimates from VBL. **D**. Accuracy component of the LME estimates from TI. The results demonstrate that TI estimates show much lower variability as compared to VBL estimates.

## Conclusion

In this paper, we have reviewed the theoretical foundation of thermodynamic integration. In the process, we have introduced the concept of free energy, which has found its way into information theory and Bayesian statistics from its origin in statistical mechanics. Approaching TI from this dual perspective allowed us to highlight the parallels and analogous concepts shared between these different scientific fields.

A key result was obtained in Eq. 23 (the TI equation), which provided (1) a graphical interpretation of the LME as the signed area under the curve given by the accuracy *A*(*β*) = −*∂F*_*H*_*/∂β*; and (2) a reliable method for estimating LME via Monte Carlo samples drawn from the power posteriors. The application of this method was demonstrated in the second part of this paper on synthetic and real-world datasets.

Specifically, we started with an experiment involving synthetic data from a linear regression model with analytical solutions for LME. This experiment demonstrated that TI produces accurate LME estimates and outperforms computationally less complex sampling-based LME estimators (AME and HME), justifying the additional complexity.

Finally, we used synthetic and real-world fMRI data to compare TI to VB, which is the current gold standard in the context of model inversion and LME estimation for DCM. Although VB was robust in most instances, we found evidence for variability in the estimates due to local optima in the objective function – especially in the case of the real-world dataset, where the model space included nonlinear DCMs, and for challenging scenarios where the number of network nodes and free parameters is high. While this problem can be ameliorated by initializing the VB algorithm from different starting points or using global optimization methods (see Lomakina et al., 2015), this would reduce computational efficiency, which is the main justification for VB as the default choice for standard applications of DCM.

Hence, sampling-based approaches like TI might become the method of choice when the robustness and validity of single-subject inference is paramount. For example, the utility of generative models for clinical applications, such as differential diagnosis based on model comparison or prediction of individual treatment responses (Stephan et al., 2017), depends on our ability to draw reliable and accurate conclusions from model-based estimates.

In addition, the experiments presented in this paper also demonstrated the practical feasibility of applying TI to complex generative models like DCM, which are characterized by high computational cost for evaluation of the likelihood function. This is made possible by an implementation that relies on parallel computing techniques, offering reasonable execution times on stand-alone workstations. Specifically, the computations for this paper were performed on a workstation equipped with an Intel Core i7 4770K (CPU) and a Nvidia Geforce GTX 1080 (GPU), with a software implementation that allow obtaining as many as 10^5^ samples of realistic DCMs in only a few minutes. Here, the important implication is that TI is no longer a method that is exclusively reserved for users with access to high performance computing clusters. The TI and DCM implementations used in this paper is available to the community as open source software (Translational Neuromodeling Unit, 2014).

## References

Annis, J., Evans, N. J., Miller, B. J., & Palmeri, T. J. (2019). Thermodynamic integration and steppingstone sampling methods for estimating Bayes factors: A tutorial. Journal of mathematical psychology, 89, 67–86. doi:10.1016/j.jmp.2019.01.005

Aponte, E. A., Raman, S., Sengupta, B., Penny, W., Stephan, K. E., & Heinzle, J. (2016). mpdcm: A toolbox for massively parallel dynamic causal modeling. Journal of Neuroscience Methods, 257, 7–16. doi:http://dx.doi.org/10.1016/j.jneumeth.2015.09.009

Bishop, C. (2006). Pattern Recognition and Machine Learning. Cambridge: Springer.

Buchel, C. (1997). Modulation of connectivity in visual pathways by attention: cortical interactions evaluated with structural equation modelling and fMRI. Cerebral cortex (New York, N.Y. 1991), 7(8), 768–778. doi:10.1093/cercor/7.8.768

Calderhead, B., & Girolami, M. (2009). Estimating Bayes factors via thermodynamic integration and population MCMC. COMPUTATIONAL STATISTICS & DATA ANALYSIS, 53, 4028–4045. doi:10.1016/j.csda.2009.07.025

Chumbley, J. R., Friston, K. J., Fearn, T., & Kiebel, S. J. (2007). A Metropolis–Hastings algorithm for dynamic causal models. NeuroImage (Orlando, Fla.), 38(3), 478–487. doi:10.1016/j.neuroimage.2007.07.028

Daunizeau, J., David, O., & Stephan, K. E. (2011). Dynamic causal modelling: A critical review of the biophysical and statistical foundations. NeuroImage, 58(2), 312–322. doi:http://dx.doi.org/10.1016/j.neuroimage.2009.11.062

David, O., Kiebel, S. J., Harrison, L. M., Mattout, J., Kilner, J. M., & Friston, K. J. (2006). Dynamic causal modeling of evoked responses in EEG and MEG. NeuroImage, 30(4), 1255–1272. doi:http://dx.doi.org/10.1016/j.neuroimage.2005.10.045

ETH Zurich. (2020). ETH Research Collection. Retrieved from https://www.research-collection.ethz.ch/bitstream/handle/20.500.11850/301664/simulation_dcms.zi p

Friston, K. J., Harrison, L., & Penny, W. (2003). Dynamic causal modelling. NeuroImage, 19(4), 1273–1302. doi:10.1016/S1053-8119(03)00202-7

Friston, K. J., Mattout, J., Trujillo-Barreto, N., Ashburner, J., & Penny, W. (2007). Variational free energy and the Laplace approximation. NeuroImage, 34(1), 220–234. doi:http://dx.doi.org/10.1016/j.neuroimage.2006.08.035

Gelman, A., & Meng, X. L. (1998). Simulating Normalizing Constants: From Importance Sampling to Bridge Sampling to Path Sampling. Statistical Science, 13(2), 163–185.

Gelman, A., & Rubin, D. B. (1992). Inference from Iterative Simulation Using Multiple Sequences. Statistical Science, 7(4), 457–472. doi:10.1214/ss/1177011136

Heinzle, J., Koopmans, P. J., den Ouden, H. E. M., Raman, S., & Stephan, K. E. (2016). A hemodynamic model for layered BOLD signals. NeuroImage (Orlando, Fla.), 125, 556–570. doi:10.1016/j.neuroimage.2015.10.025

Jaynes, E. T. (1957). Information Theory and Statistical Mechanics. Physical Review, 106(4), 620–630. doi:10.1103/physrev.106.620

Kirkwood, J. G. (1935). Statistical Mechanics of Fluid Mixtures. The Journal of chemical physics, 3(5), 300–313. doi:10.1063/1.1749657

Landau, D. P. (2015). A guide to monte carlo simulations in statistical physics (4th ed. ed.): Cambridge : University Press.

Lartillot, N., & Philippe, H. (2006). Computing Bayes Factors Using Thermodynamic Integration. Systematic Biology, 55(2), 195–207. doi:10.1080/10635150500433722

Lomakina, E. I., Paliwal, S., Diaconescu, A. O., Brodersen, K. H., Aponte, E. A., Buhmann, J. M., & Stephan, K. E. (2015). Inversion of hierarchical Bayesian models using Gaussian processes. NeuroImage, 118, 133–145. doi:http://dx.doi.org/10.1016/j.neuroimage.2015.05.084

MacKay, D. J. C. (2004). Information Theory, Inference, and Learning Algorithms (Repr. with corr. ed.). Cambridge: Univ. Press.

Marreiros, A. C., Kiebel, S. J., & Friston, K. J. (2008). Dynamic causal modelling for fMRI: A two-state model. NeuroImage (Orlando, Fla.), 39(1), 269–278. doi:10.1016/j.neuroimage.2007.08.019

McDowell, J. E., Dyckman, K. A., Austin, B. P., & Clementz, B. A. (2008). Neurophysiology and neuroanatomy of reflexive and volitional saccades: Evidence from studies of humans. Brain and cognition, 68(3), 255–270. doi:10.1016/j.bandc.2008.08.016

Moran, R., Pinotsis, D. A., & Friston, K. (2013). Neural masses and fields in dynamic causal modeling. Frontiers in computational neuroscience, 7, 57–57. doi:10.3389/fncom.2013.00057

Neal, R. M., & Hinton, G. E. (1998). A View of the Em Algorithm that Justifies Incremental, Sparse, and other Variants. In M. I. Jordan (Ed.), Learning in Graphical Models (pp. 355–368). Dordrecht: Springer Netherlands.

Ortega, P. A., & Braun, D. A. (2013). Thermodynamics as a theory of decision-making with information-processing costs. Proceedings of the Royal Society. A, Mathematical, physical, and engineering sciences, 469(2153), 20120683. doi:10.1098/rspa.2012.0683

Penny, W., & Sengupta, B. (2016). Annealed Importance Sampling for Neural Mass Models. PLOS Computational Biology, 12(3), e1004797–e1004797. doi:10.1371/journal.pcbi.1004797

Penny, W., Stephan, K. E., Mechelli, A., & Friston, K. J. (2004a). Comparing dynamic causal models. NeuroImage, 22(3), 1157–1172. doi:http://dx.doi.org/10.1016/j.neuroimage.2004.03.026

Penny, W., Stephan, K. E., Mechelli, A., & Friston, K. J. (2004b). Modelling functional integration: a comparison of structural equation and dynamic causal models. NeuroImage, 23, S264–S274. doi:https://doi.org/10.1016/j.neuroimage.2004.07.041

Raman, S., Deserno, L., Schlagenhauf, F., & Stephan, K. E. (2016). A hierarchical model for integrating unsupervised generative embedding and empirical Bayes. Journal of Neuroscience Methods, 269, 6–20. doi:http://dx.doi.org/10.1016/j.jneumeth.2016.04.022

Schwarz, G. (1978). Estimating the Dimension of a Model. The Annals of statistics, 6(2), 461–464. doi:10.1214/aos/1176344136

Sengupta, B., Friston, K. J., & Penny, W. (2015). Gradient-free MCMC methods for dynamic causal modelling. NeuroImage (Orlando, Fla.), 112(C), 375–381. doi:10.1016/j.neuroimage.2015.03.008

Sengupta, B., Friston, K. J., & Penny, W. (2016). Gradient-based MCMC samplers for dynamic causal modelling. NeuroImage (Orlando, Fla.), 125, 1107–1118. doi:10.1016/j.neuroimage.2015.07.043

Stephan, K. E., Kasper, L., Harrison, L. M., Daunizeau, J., den Ouden, H. E. M., Breakspear, M., & Friston, K. J. (2008). Nonlinear dynamic causal models for fMRI. NeuroImage, 42(2), 649–662. doi:http://dx.doi.org/10.1016/j.neuroimage.2008.04.262

Stephan, K. E., Penny, W., Daunizeau, J., Moran, R. J., & Friston, K. J. (2009). Bayesian model selection for group studies. NeuroImage, 46(4), 1004–1017. doi:http://dx.doi.org/10.1016/j.neuroimage.2009.03.025

Stephan, K. E., Schlagenhauf, F., Huys, Q. J. M., Raman, S., Aponte, E. A., Brodersen, K. H., … Heinz, A. (2017). Computational neuroimaging strategies for single patient predictions. NeuroImage, 145, Part B, 180–199. doi:https://doi.org/10.1016/j.neuroimage.2016.06.038

Swendsen, R. H., & Wang, J.-S. (1986). Replica Monte Carlo Simulation of Spin-Glasses. Physical Review Letters, 57(21), 2607–2609. doi:10.1103/physrevlett.57.2607

Translational Neuromodeling Unit. (2014). TAPAS - Translational Algorithms for Psychiatry-Advancing Science. Retrieved from http://www.translationalneuromodeling.org/tapas

Watanabe, S. (2013). A Widely Applicable Bayesian Information Criterion. Journal of Machine Learning Research, 14, 867–897.

Welvaert, M., & Rosseel, Y. (2013). On the Definition of Signal-To-Noise Ratio and Contrast-To-Noise Ratio for fMRI Data. PLOS ONE, 8(11), e77089. doi:10.1371/journal.pone.0077089

Wipf, D., & Nagarajan, S. (2009). A unified Bayesian framework for MEG/EEG source imaging. NeuroImage, 44(3), 947–966. doi:https://doi.org/10.1016/j.neuroimage.2008.02.059

Yao, Y., Raman, S. S., Schiek, M., Leff, A., Frässle, S., & Stephan, K. E. (2018). Variational Bayesian inversion for hierarchical unsupervised generative embedding (HUGE). NeuroImage, 179, 604–619. doi:https://doi.org/10.1016/j.neuroimage.2018.06.073

Yao, Y., & Stephan, K. E. (2020). Markov chain Monte Carlo methods for hierarchical clustering of dynamic causal models. Retrieved from https://arxiv.org/abs/2012.05744

